# *BridgePRS*: A powerful trans-ancestry Polygenic Risk Score method

**DOI:** 10.1101/2023.02.17.528938

**Authors:** Clive Hoggart, Shing Wan Choi, Judit García-González, Tade Souaiaia, Michael Preuss, Paul O’Reilly

**Affiliations:** Department of Genetics and Genomic Sciences, Icahn School of Medicine, Mount Sinai, 1 Gustave L. Levy Pl, New York, 10029, NY, USA; Department of Cellular Biology, Suny Downstate Health Sciences, Brooklyn, NY, USA; The Charles Bronfman Institute for Personalized Medicine, Icahn School of Medicine, Mount Sinai, 1 Gustave L. Levy Pl, New York, 10029, NY, USA

## Abstract

Polygenic Risk Scores (PRS) have huge potential to contribute to biomedical research and to a future of precision medicine, but to date their calculation relies largely on European-ancestry GWAS data. This global bias makes most PRS substantially less accurate in individuals of non-European ancestry. Here we present *BridgePRS*, a novel Bayesian PRS method that leverages shared genetic effects across ancestries to increase the accuracy of PRS in non-European populations. The performance of *BridgePRS* is evaluated in simulated data and real UK Biobank (UKB) data across 19 traits in African, South Asian and East Asian ancestry individuals, using both UKB and Biobank Japan GWAS summary statistics. *BridgePRS* is compared to the leading alternative, *PRS-CSx*, and two single-ancestry PRS methods adapted for trans-ancestry prediction. PRS trained in the UK Biobank are then validated out-of-cohort in the independent Mount Sinai (New York) Bio*Me* Biobank. Simulations reveal that *BridgePRS* performance, relative to *PRS-CSx*, increases as uncertainty increases: with lower heritability, higher polygenicity, greater between-population genetic diversity, and when causal variants are not present in the data. Our simulation results are consistent with real data analyses in which *BridgePRS* has better predictive accuracy in African ancestry samples, especially in out-of-cohort prediction (into Bio*Me*), which shows a 60% boost in mean *R*^*2*^ compared to *PRS-CSx* (*P* = 2 *×* 10^*−6*^). *BridgePRS* performs the full PRS analysis pipeline, is computationally efficient, and is a powerful method for deriving PRS in diverse and under-represented ancestry populations.

## Introduction

Polygenic Risk Scores (PRS) are mostly derived using European ancestry genome-wide association study (GWAS) data, which results in substantially lower predictive power when applied to non-European samples, in particular African ancestry samples [1, 2]. The PRS trans-ancestry portability problem is well-established and is due to marked linkage disequilibrium (LD) differences, allele frequency differences driven by genetic drift and natural selection, and GxE interactions affecting causal effect sizes [3]. Consequently, the aetiological insights and clinical utility provided by PRS derived in Europeans may have limited relevance to individuals of non-European ancestries.

Increasing GWAS sample sizes of underrepresented populations will help to improve their PRS, but optimal power will be achieved by utilising all GWAS available across ancestries for PRS prediction into any one ancestry. This is the approach of *PRS-CSx* [4], developed to tackle the PRS portability problem, which makes cross-population inference on the inclusion of each SNP across the genome (or more precisely, the degree of shrinkage of variant effect sizes to zero). *PRS-CSx* utilises Bayesian modelling with a prior that strongly shrinks small effect sizes to zero, reducing the number of candidate SNPs to a minimal set. This is analogous to fine-mapping of causal variants. However, while the inclusion of causal variants in the PRS is ideal, fine-mapping approaches may not be as effective when causal variants are missing or are underpowered to be identified.

We introduce *BridgePRS*, a novel Bayesian PRS method that also integrates trans-ancestry GWAS summary statistics. Unlike the fine-mapping approach of *PRS-CSx, BridgePRS* aggregates information across putative loci by estimating optimal SNP weights to best tag causal variants. The focus is on correctly estimating effect sizes, rather than location, which is key when prediction is the goal. This approach is less reliant on the inclusion of causal variants. *BridgePRS* is most applicable to combining the information of a well-powered GWAS performed in a (discovery) population(s) not matched to the ancestry of the target sample, with a second GWAS of limited power in a (target) population that is well-matched to the ancestry of the target sample. We apply *BridgePRS* to simulated data and compare its performance to *PRS-CSx* and two single ancestry PRS methods adapated to use trans-ancestry GWAS data. The simulations demonstrate the different scenarios in which *BridgePRS* and *PRS-CSx* are optimal. We then utilise UK Biobank (UKB) [5] and Biobank Japan (BBJ) [6, 7] GWAS data to construct PRS for African, South Asian and East Asian ancestry samples. Resultant PRSs are then validated in the UKB and in the entirely independent Mount Sinai Bio*Me* Biobank (Bio*Me*) [8], producing results consistent with the simulations.

## Results

### Overview of *BridgePRS* method

An overview of the *BridgePRS* modelling employed is shown in Figure 1. The key modelling (see Methods) is broken into two stages: (1) a PRS is trained and optimised using discovery population (e.g. European) data, with a zero-centred Gaussian prior distribution for SNP effect sizes (analogous to ridge regression) within putative loci, (2) the SNP effect sizes of this PRS are treated as priors and updated in a Bayesian framework by those of the target population (e.g. African) GWAS. Thus, this two-stage <u>B</u>ayesian-<u>ridge</u> approach of *BridgePRS*, “bridges” the PRS between the two populations.

**Fig. 1.**
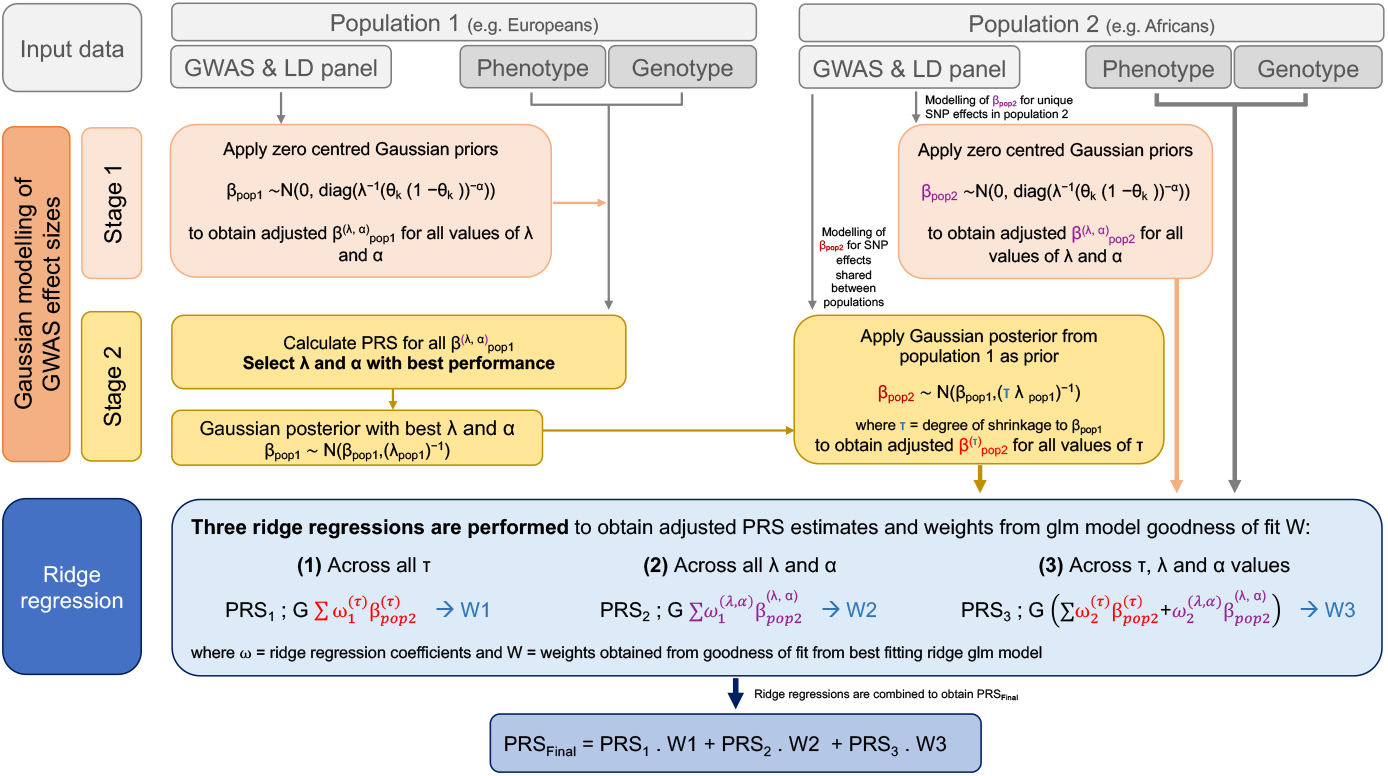
Flow diagram describing the modelling of BridgePRS.

The main causes of poor trans-ancestry PRS portability are differences in LD and allele frequencies between populations [3]. Differences in LD result in the best tag for a causal variant differing between populations. To account for the resultant uncertainty in the location of causal variants, *BridgePRS* averages SNP effects across putative loci, instead of selecting a single best SNP as performed by standard clumping+thresholding (C+T) PRS [9]. *BridgePRS* is first applied to the discovery population GWAS, using Bayesian modelling with zero-centred Gaussian priors, equivalent to penalised likelihood ridge regression, at putative loci. Given summary data from large GWASs in Europeans, we find that this procedure alone significantly improves predictive accuracy in African and South Asian target data compared to choosing single best SNPs at putative loci.

This first stage modelling results in multivariate Gaussian posterior distributions for SNP effect sizes at each locus. In the next stage, *BridgePRS* integrates the (smaller) target population GWAS data into the PRS by using this posterior distribution as a prior distribution for SNP effect sizes of the target population. This stage allows for different effect size estimates between the populations, accounting for differences in allele frequencies driven by drift or selection, LD and GxE interactions affecting causal effect sizes. Both stages of the modelling use conjugate prior-posterior updates, providing computationally efficient analytical solutions and enabling BridgePRS analyses to be performed rapidly.

Variation in causal allele frequencies between populations can mean that causal variants with relatively low minor allele frequency in the discovery population are estimated with large errors or are missed altogether. To ameliorate this problem, PRS are derived by applying *BridgePRS* stage 1 modelling to the target population data alone (see Methods).

Each stage of the modelling is fit across a spectrum of prior parameters and criteria to select loci for inclusion in the PRS calculation, with each combination of parameters giving rise to a unique PRS. These PRS are then combined in a ridge regression fit using available genotype-phenotype test data, choosing the optimum ridge penalty parameter by cross-validation (see Methods).

### Benchmarking methods via simulation

We used the HAPGEN2 software [10] to simulate HAPMAP3 variants for 100K Europeans, 40K Africans and 40K East Asian ancestry samples using 1000G Phase 3 samples [11] as reference. Simulations were restricted to 1,295,289 variants with minor allele frequency *>* 1% in at least one of the three populations. Phenotypes were subsequently simulated under three models of genetic architecture in which causal variants were sampled from 1%, 5% and 10% of the available HAPMAP3 variants. Population-specific effect sizes were sampled from a multivariate Gaussian distribution with between-population correlation of 0.9. Genetic effects were combined assuming additivity and Gaussian noise at two levels of variance were added to generate phenotypes with 25% and 50% SNP-heritability. For each of the six scenarios of polygenicity and heritability, ten independent phenotypes were generated and analyses were run with and without including the causal variants.

Data were split into training for GWASs (80K European, 20K non-European) with the remainder split equally into 10K samples for model optimisation (test data) and assessment of model performance (validation data). The performance of *BridgePRS* was compared with *PRS-CSx*, an alternative trans-ancestry PRS method, *PRS-CS-mult* and *PRSice-meta. PRS-CS-mult* applies the single ancestry *PRS-CS* method [12] to the populations under study and combines them by estimating weights in a linear regression utilising the test data. *PRSice-meta* applies clumping and thresholding, as implemented in *PRSice* [13], to the meta-analysis of the populations under study using reference LD panels from the same populations and selecting the PRS that optimises prediction in the test data.

Polygenicity varying between 1%-10% (fraction of variants with non-zero effect sizes) is consistent with a recent study of 28 complex traits in the UK Biobank [14]. Between-population correlation of causal variant effect sizes of 0.9 is consistent with a recent multi-ancestry lipids GWAS in which causal variants were fine-mapped [15]. Approximately one-third to two-thirds of heritability is captured by common SNPs [16] and, therefore, our simulation at 25% heritability implies a total heritability 37.5% - 75%. Power of GWAS, and therefore PRS, is a function of sample size and heritability, such that doubling heritability is equivalent to doubling sample size in terms of power. Therefore, our simulations at 50% heritability and GWASs with 80K European samples are equivalent to 25% heritability and GWASs with 160K European samples.

Figure 2 summaries the results from PRS analyses performed on simulated data. Both *BridgePRS* and *PRS-CSx* outperform the single ancestry methods across all scenarios. *BridgePRS* outperforms *PRS-CSx* in all analyses of African samples with 5% and 10% of variants assigned as causal. With 1% of variants causal the methods have similar accuracy when causal variants are not included and *PRS-CSx* performs better with causal variants included. In analyses of East Asian samples, the same relative pattern is observed, but the differences are less pronounced and *PRS-CSx* performs better in all scenarios in which 1% of variants are causal. Across all analyses, *BridgePRS* performs relatively better compared to *PRS-CSx* when the causal variants are not included in the data (Supplementary Figure 1). Overall, the simulations reveal that *BridgePRS* performance, relative to *PRS-CSx*, increases as uncertainty increases: at lower heritability, higher polygenicity, greater between-population genetic diversity, and when causal variants are not present in the data.

**Fig. 2.**
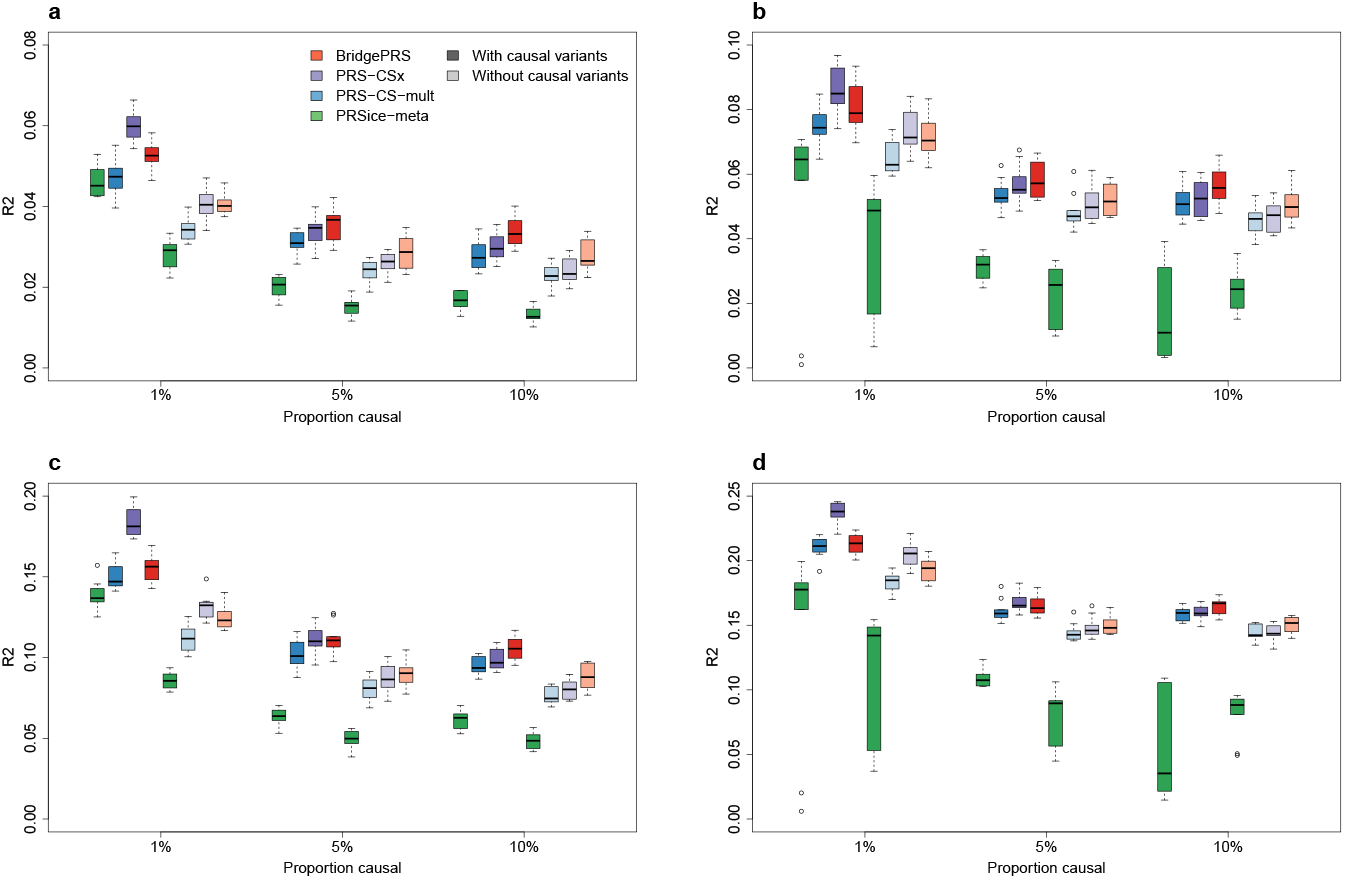
Phenotypic variance explained by *BridgePRS, PRS-CSx, PRS-CS-mult* and *PRSicemeta* across six siumulation scenarios, with and without the causal variants included in the model, ten simulated phenotypes per scenerio. **a** African ancestry samples for phenotypes with 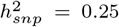 b East Asian ancestry samples for phenotypes with 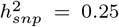 African ancestry samples for phenotypes with 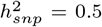 d East Asian ancestry samples for phenotypes with 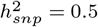

These analyses used 1000G data as their reference LD panel, i.e. the correct LD panel. To assess the sensitivity of the methods to misspecification of LD, analyses were rerun using UKB data to estimate ancestry-specific LD. Supplementary Figure 2 shows the performance of *BridgePRS* and *PRS-CSx* using an LD reference panel constructed from African and East Asian UKB samples relative to their performance using the 1000G reference panel. Both methods exhibit a minimal loss in predictive accuracy using UKB reference panels.

### Benchmarking methods via real data: UK Biobank and Bio*Me* Biobank

The four PRS methods were applied to UK Biobank (UKB) [5] samples of African and South Asian ancestry across 19 continuous biometric and biochemical traits (for East Asian ancestry see below). These traits were selected to maximise heritability and samples sizes of non-European individuals and to minimise their pairwise correlation (maximum *r*^2^ < 0.3; see Methods). For each trait, UKB samples of European, African and South Asian ancestry were split into training, test and validation sets in proportions of 2/3, 1/6 and 1/6, respectively. Sample sizes are shown in Supplementary Table 1. The training data were used to generate GWAS summary statistics and the test data used to select optimal model parameters. Results are shown for the resultant PRS in the unseen UKB validation data. In addition, an entirely out-of-sample validation study was performed by applying the PRS derived in the UKB to the Mount Sinai Bio*Me* Biobank (Bio*Me*) [8] for the nine traits also available in Bio*Me*.

Within the UKB there are 2,472 East Asian samples, which is too few to split into training (GWAS), test and validation sets as above. However, GWAS summary statistic data from Biobank Japan (BBJ) are available for download [6, 7]. We combined these data with the European UKB GWAS summary statistics described above for ten overlapping traits to estimate PRS for East Asians ancestry (as above). *BridgePRS* combines SNP effect size estimates across GWASs (as does the *PRSice-meta* method) and, therefore, requires effect sizes to be on the same scale. However, the BBJ summary statistics were generated after standardising the trait values to have mean zero and standard deviation of one, whereas the UKB GWASs were applied to raw trait data. Therefore, before applying the methods, the BBJ effect estimates and standard errors were transformed to the respective scale of the UKB measures assuming that the BBJ and UKB trait values had the same variance. UKB East Asian samples were then split equally into test data for model optimisation and validation data to assess model performance, as above. PRS were also validated in East Asian Bio*Me* samples across eight overlapping traits.

Trait sample sizes for each ancestral population in the UKB and Bio*Me* cohorts are shown in Supplementary Tables 1 and 2. For all analyses, imputed genotype data were used. Figure 3 shows the PRS variance explained (*R*^2^ with CIs; see Methods) by *BridgePRS, PRS-CSx, PRS-CS-mult* and *PRSice-meta*, averaged across all traits, for prediction into African, South Asian and East Asian ancestry samples in the UKB and Bio*Me* cohorts. Also shown are *P* -values comparing *R*^2^ between *BridgePRS, PRS-CSx* and *PRS-CS-mult* (not *PRSice-meta* since it is universally inferior across all comparisons). For prediction into African ancestry samples, *BridgePRS* has the highest average *R*^2^ in both cohorts, significantly so (*P* =2×10^−6^ Vs *PRS-CSx*) for the out-of-cohort prediction into Bio*Me* with an average relative boost in *R*^2^ of 60%. For prediction into South Asian ancestry there are no significant differences between methods. For prediction into East Asians, *BridgePRS* is inferior to both *PRS-CSx* and *PRS-CS-mult* in both UKB and Bio*Me*, but only significantly so within-cohort in the UKB, and there are no significant differences between *PRS-CSx* and *PRS-CS-mult* in both cohorts.

**Fig. 3.**
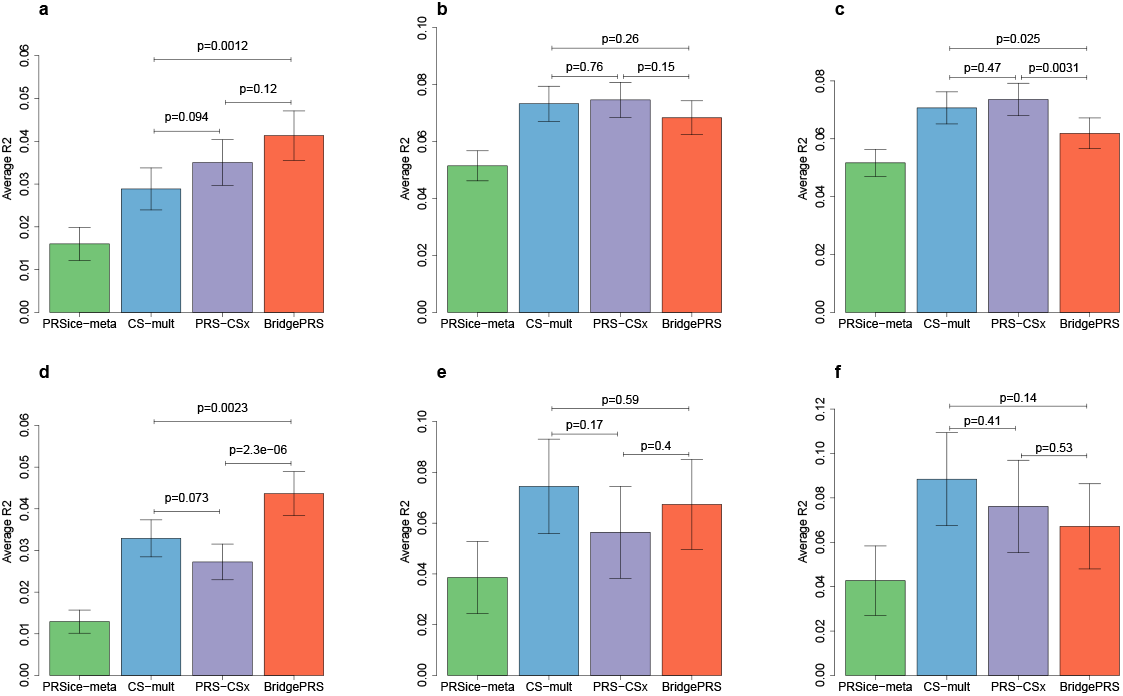
Average phenotypic variance explained by *BridgePRS, PRS-CSx, PRS-CS-mult* and *PRSice-meta* in samples of African, South Asian and East Asian ancestry in the UK Biobank and Bio*Me* cohorts. **a** African ancestry samples in UK Biobank. **b** South Asian ancestry samples in UK Biobank. **c** East Asian ancestry samples in UK Biobank. **d** African ancestry samples in Bio*Me*. **e** South Asian ancestry samples in Bio*Me*. **f** East Asian ancestry samples in Bio*Me*.

Figure 4 shows the individual results for each trait (*R*^2^ with CIs) analysed in the out-of-sample prediction into the Bio*Me* cohort. While the methods show similar results across many of the traits, the relative performance of the methods is highly variable and for some traits there are distinct differences in accuracy of the methods, especially in African ancestry samples. For example, in African ancestry samples, *BridgePRS* performs markedly better for mean corpuscular volume (MCV) and LDL, but markedly worse for Eosinophil count (Eos). In both African and South Asian ancestry, the *PRS-CSx* prediction of height is highly inaccurate, which may be due to the impact of variant overlap between cohorts when applying *PRS-CSx* out-of-sample (see Discussion). The corresponding trait-specific results for prediction into UKB are shown in Supplementary Figures 3 and 4, with a similar pattern of results observed. Of note, *BridgePRS* again performs markedly better for mean corpuscular volume and LDL in African ancestry samples.

**Fig. 4.**
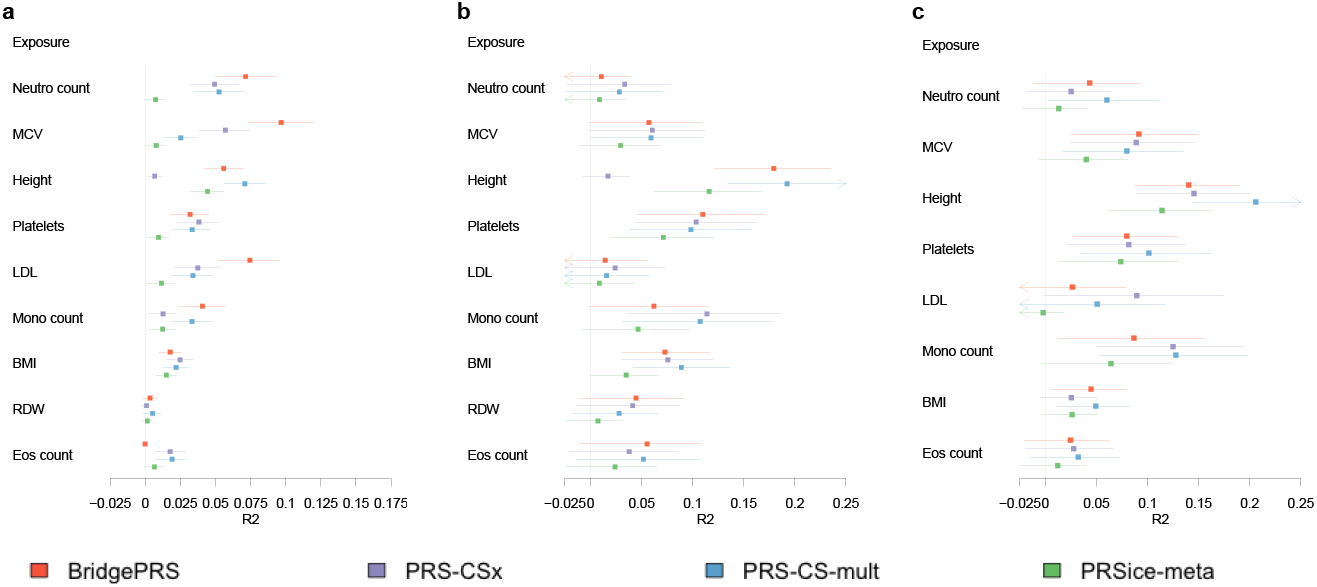
Phenotypic variance explained (*R*2 point estimates and 95% confidence intervals) by *BridgePRS, PRS-CSx, PRS-CS-mult* and *PRSice-meta* in samples of African, South Asian and East Asian ancestry in the Bio*Me* cohort. **a** African ancestry samples. **b** South Asian ancestry samples. **c** East Asian ancestry samples. Neutro count=Neutrophil count, MCV=Mean corpuscular volume, Platelets=Platelet count, Mono count=Monocyte count, BMI=Body mass index, RDW=Red blood cell distribution width, Eos count=Eosinophil count.

## Discussion

We have introduced a novel trans-ancestry PRS method, *BridgePRS*, that leverages shared genetic effects across ancestries to increase the accuracy of PRS in non-European populations. We benchmarked *BridgePRS* and leading trans-ancestry PRS method *PRS-CSx*, as well as single ancestry PRS methods *PRS-CS* and *PRSice* adapted for trans-ancestry prediction, across a range of simulated and real data. In all analyses, target population PRS utilise GWAS summary statistics from Europeans and the target population. Results from our simulated data reveal that *BridgePRS* has higher performance relative to *PRS-CSx* when uncertainty is greater: for lower heritability traits, for lower GWAS sample sizes (especially for the target population), when genetic signal is dispersed over more causal variants (higher polygenicity), for greater between-population diversity (e.g. with European base and African target, rather than Asian target), and when the causal variants are not included in the analyses. In all analyses of simulated data, *BridgePRS* and *PRS-CSx* had superior performance relative to the single-ancestry PRS methods.

Application of the methods to real GWAS summary statistics from the UK Biobank (UKB) and Biobank Japan (BBJ) and validation in independent samples of African, South Asian and East Asian ancestry in UKB and Bio*Me* (recruited in the New York City area of the USA) gave results consistent with the simulations. Specifically, *BridgePRS* has superior average *R*^2^ across the traits analysed for samples of African ancestry in which uncertainty is high due to greater differences in LD between Africans and Europeans, and because of the relatively small African GWASs used. Likewise, *PRS-CSx* has superior average *R*^2^ for samples of East Asian ancestry for which differences in LD are smaller and the contributing East Asian GWASs are much larger (90K-160K). For prediction into South Asian ancestry, in which LD is relatively similar, but the South Asian GWASs used are small, the methods perform similarly.

The stronger performance of *PRS-CSx* in the real data analysis of East Asians may also be due to *PRS-CSx* not requiring GWASs to be on the same scale and, thus, being unaffected by the rescaling of the BBJ effect estimates. *PRS-CSx* is unaffected by GWAS scale as it combines information across ancestries on the shrinkage (to zero) of the effect estimate of each SNP and does not combine information on effect sizes. The final *PRS-CSx* PRS estimate is derived by combining ancestry-specific PRSs with relative weights estimated in a linear regression in the test data. Differences in scale between the base GWASs will be accounted for by the linear regression weights. *BridgePRS* should have improved performance when the GWASs used are performed on the same scale, since it shares information on effect sizes across ancestries.

Using UK Biobank and Bio*Me* data, we have demonstrated that *BridgePRS* has superior out-of-cohort predictive accuracy in genetic prediction in individuals of African ancestry. However, *PRS-CSx* has better accuracy when using UKB European and BBJ East Asian summary statistics to predict into individuals of East Asian ancestry. In general, in simulated and real data, *BridgePRS* performs better than *PRS-CSx* when uncertainty in mapping of causal variants is higher. Given the complementary nature of the two methods, either can be optimal depending on the trait and study characteristics, and therefore we recommend applying both methods until it is known which offers greater power in the given setting.

*BridgePRS* is a fully dedicated PRS tool that performs the entire PRS process and offers a novel theoretical approach to tackling the PRS portability problem, with particularly strong performance for deriving PRS in African and other diverse and under-represented ancestry populations.

### Software availability

Software implementing *BridgePRS* with documentation and example data can be downloaded from GitHub (https://github.com/clivehoggart/BridgePRS).

## Acknowledgements

We thank the participants in the UK Biobank and the scientists involved in the construction of this resource and Biobank Japan (BBJ) for releasing the genome-wide association summary. This research has been conducted using the UK Biobank Resource under application 18177 (Dr O’Reilly). All participants gave full informed consent. This work was supported by a grant from the National Institute of Health (R01MH122866) to PFO and through the computational resources and staff expertise provided by Scientific Computing at the Icahn School of Medicine at Mount Sinai.

## Methods

### The *BridgePRS* model

All modelling is performed at the locus level and each locus is assumed to be independent of all others. Within loci, SNP effect sizes *β* are modelled by a multivariate Gaussian distribution and we assume the trait, *y*, of individuals with genotype data, X, at the locus follows a Gaussian distribution *y ∼* N(*Xβ, ψI*). Throughout, the Gaussian distribution is parameterised by its mean and precision matrix (=inverse covariance matrix).

Below we describe *BridgePRS* methodology to derive a PRS for a target population, population 2, (in our applications: African, South Asian and East Asian) for which we have summary statistics from a relatively under-powered GWAS and GWAS summary statistics from a well powered GWAS from a different ancestral population, population 1 (in our application: European). We also assume we have small data sets of genotype-phenotype data from both populations.

### Stage 1: PRS informed by a single population

In stage 1 modelling, we train and optimise PRS using GWAS summary statistics and test genotype-phenotype data from a single population. To determine the PRS for population 2, this modelling stage is applied to populations 1 and 2 independently. Application to population 1 determines the prior distributions for population 2 SNP effects used in stage 2, see below, and application to population 2 is used to identify population 2 specific effects and thus missed in population 1.

In stage 1, a zero-centred conjugate Gaussian prior is assigned for the SNP effects at each locus *β ∼*N(0, *ψλI*), where *I* is the identity matrix and *λ* is a vector of SNP specific shrinkage parameters. The use of a conjugate prior allows the posterior distribution of SNP effects to be determined analytically [18]:

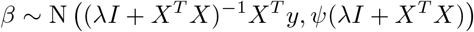

*X*^*T*^ *y* can be calculated from the vector of maximum likelihood marginal effects 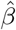 available from GWAS summary statistics by 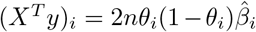 where *n* is the sample size, *θ* is the vector of allele frequencies (AF) and (*X*^*T*^ *y*)_*i*_ is the *i*^*th*^ element of *X*^*T*^ *y, i* indexes SNPs. *X*^*T*^ *X* = *n*Φ, where Φ is the pairwise genotypic covariance which can be estimated from a reference panel representative of the population used in the GWAS. Thus, rescaling *λ* by *n*, the posterior is estimated as

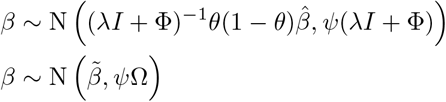

To accommodate the effects of natural selection, we allow the prior on SNP effects to be dependent on AF such that the prior precision for the *k*^*th*^ SNP is *λ*_*k*_ = *λ*_0_(*θ*_*k*_(1 − *θ*_*k*_))^*α*^ and *α∈* (0, 1) [19]. When *α* = 0 AF and effect size are *a priori* independent, *α* = 1 is the value implicitly assumed by many methods [20], which implies a strong assumption of larger effects at lower minor allele frequency SNPs. Multiple models are fit at each locus under priors defined by all combinations of *α* = (0, 0.25, 0.5, 0.75, 1) and *λ*_0_ = (0.05, 0.1, 0.2, 0.5, 1, 2, 5). Loci are ranked by the *P* -value of their most associated SNP and assigned to subset *S*_*k*_ if the top SNP *P* -value < 10^*−k*^, values of *k* = 1, …, 8 are considered. Multiple genome-wide PRS are calculated for a test set of phenotype and genotype data by summing the effects across all contributing loci across all combinations of *α, λ*_0_ and *k*:

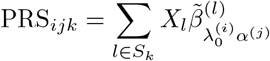

where *X*_*l*_ is the genotype data at locus *l*, 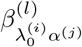 is the posterior mean at locus *l* with prior defined by parameters 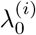 and *α*^(*j*)^, and *S*_*k*_ is the subset of loci with top SNP *P* -value < 10^−*k*^. A single PRS is calculated by a weighted sum of the PRS across all *i, j* and *k*, with weights determined by a ridge regression fit to the test data utilising leave-one-out cross-validation to select the ridge shrinkage parameter, which minimises out-of-sample deviance as implemented in the R package glmnet [21].

### Stage 2: PRS informed by stage 1

In stage 2 modelling, SNP effect sizes estimated by the application of stage 1 modelling to population 1 (eg. Europeans) are updated based on population 2 GWAS summary statistics and optimised using population 2 genotypephenotype data. The prior used is taken as the posterior derived from the *λ*_0_ and *α* prior parameters which optimise prediction in the test data of population

1. As for stage 1, this prior is also a multivariate Gaussian. A parameter *τ* is added to the precision parameter of the Gaussian to control the contribution of population 1 to population 2, thus the prior is specified as 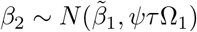. This is similarly a conjugate model with a Gaussian posterior [18]:

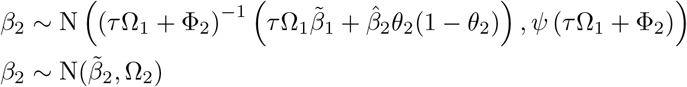

where Φ_2_ is the SNP covariance at the locus in population 2, 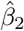 is the vector of marginal maximum likelihood SNP effect sizes and *θ*_2_ is the vector of allele frequencies. Small values of *τ* correspond to using effect estimates close to those from population 2, as *τ* increases more weight is assigned to population 1, such that as *τ → ∞, β*_2_ *→ β*_1_.

### Ranking Loci in Stage 2

Because of differences in LD between populations, we do not rank loci by the *P* -value of a single best SNP, we aggregate information across loci by adapting the F-test. It can be shown that the F-test in a multivariate linear regression model for the null *H*_0_: *β* = 0, is well approximated by (see Supplementary Material for proof):

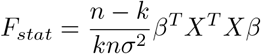

with degrees of freedom *k* and *n* − *k*, where *k* is the dimension of of *β, n* is the number of observations and *σ*^2^ is the phenotypic variance. The maximum likelihood estimate and *X*^*T*^ *X* are substituted by the posterior mean and precision matrix and *n* with *n*_*eff*_ = *n*(1 + *τ*), the effective number of observations accounting for the prior giving the statistic

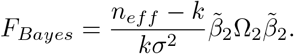

The resultant tail probability is analogous to a *P* -value, although the parameter estimates *β* and *λ* include prior information and, thus, it cannot be interpreted as such. Instead, for each *τ*, a locus with test statistic F is assigned to *S*_*k*_ if *F > q*_*k*_, where *q*_*k*_ is the F quantile corresponding to Prob(*p <* 10^*−k*^), where *p* are the locus-specific top SNP *P* -values. This ranking ensures the pseudo F-statistic ranking assigns the same number of loci to each subset as the SNP *P* -value ranking. As for the one stage single-ancestry PRS, multiple genome-wide PRS are constructed by:

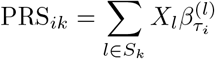

where 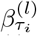 is the posterior mean at locus *l* with prior defined by parameters *τ* ^(*i*)^, and *S*_*k*_ is the subset of loci with *F* > *q*_*k*_. Models are fit for *τ* = 1, 2, 5, 10, 15, 20, 50, 100, 200, 500 and the same *P* -value thresholds as used in the first stage of the modelling. A single PRS is estimated via a ridge regression fit using population 2 test data as described above using glmnet.

The average *R*^2^ achieved by ranking loci by the pseudo F-statistic versus *P* - value from the European GWASs across the 19 traits analysed in this paper for African and South Asian UKB sample are 0.0413 v 0.0403 and 0.0683 v 0.0688 respectively, while we observed a more pronounced improvement in analyses in genotyped data (not shown here) of: 0.0413 v 0.0359 and 0.0694 v 0.0646, with *P* -value for superiority of F-statistic ranking 0.086. All results presented used the pseudo F-statistic loci ranking. However, the *BridgePRS* script allows users to rank loci in stage 2 using either ranking methods.

### Incomplete SNP overlap between populations 1 and 2

Quality control (QC) is performed separately in each population, see below. This results in variants included in analyses differing between populations. Thus, stage 2 analyses are performed on the intersection of variants passing QC in both populations and the prior is calculated conditional on effects of non-overlapping variants set to zero. Thus, given a prior of 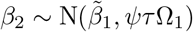, the prior on the overlapping variants is given by [18]

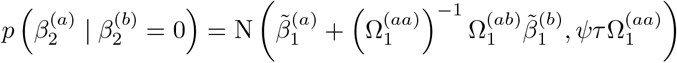

where *a* represents the overlapping variants, *b* the non-overlapping variants and 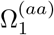 and 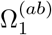 are the appropriate submatrices of Ω_1_. SNP overlap is taken at stage 2 to allow models fit in stage 1 to be applied to other data sets with different SNP sets.

### Combining PRS

We consider three alternative models for the PRS of population 2: (1) PRS estimated using only population 2, i.e. European GWAS does not inform the PRS of population 2, (2) PRS estimated using only the two-stage European informed PRS, i.e. the population 2 GWAS is under-powered and contributes insufficient information on its own and (3) both the population 2 only PRS and the two-stage PRS contribute independent information. The estimation of models (1) and (2) are determined by a cross-validated ridge regression fit as described above using glmnet. Model (3) is estimated similarly by merging all single ancestry and two stage PRS and weighting by a cross-validated ridge regression fit.

The final PRS is a weighted average of these three PRS, with weights determined by the estimated marginal likelihood of each. The log-marginal likelihood of a linear regression model *M*_*i*_ can be approximated by [22]

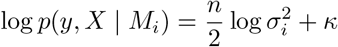

where 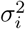 is the residual model variance estimated from cross-validation and *κ* is a constant. With equal prior weight for each of the models, the posterior model weights for models *M*_1_, *M*_2_ and *M*_3_ are given by:

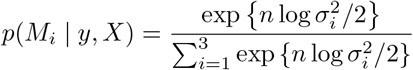

Combining PRS in this way can be extended to any number of contributing PRS. For example, we additionally combined PRS for African ancestry samples constructed from East Asian BBJ and African UKB GWAS summary statistics to PRS constructed in our main analysis which used African and European UKB GWAS summary statistics. Supplementary Figure 5 compares trait *R*^2^ for AFR+EUR PRS with AFR+EUR+EAS PRS for UKB and BBJ overlapping traits. Marginal improvement is observed by the addition of the BBJ East Asian data, for monocyte count, BMI and height, for the other traits *R*^2^ is practically unaltered.

### Definition of loci

Loci for the two-stage modelling were defined by clumping and thresholding of European GWAS summary statistics and LD structure using PLINK v1.9 [23] with the following parameters: --clump-p1 0.01 --clump-p2 0.01 --clump-kb 1000 --clump-r2 0.01. The *P* -value for each locus was determined by the *P* -value of the lead SNP of the locus in the European GWAS. The ancestry specific loci were defined similarly but used GWAS data from the appropriate ancestry.

### Application of *PRS-CSx*

*PRS-CSx* is a python based software package that integrates GWAS summary statistics and LD reference data from multiple populations to estimate population specific PRS. *PRS-CSx* applies a continuous shrinkage prior to SNP effects genome-wide in which the sparseness of the genetic architecture across populations is controlled by a parameter ϕ. PRS-CSx does not make inference on ϕ but instead estimates separate PRS for each value of *ϕ* considered. Throughout we follow the implementation described in Ruan et al [4], thus values of *ϕ* = (10^−6^, 10^−4^, 10^−2^, 1) were considered. For each *ϕ, PRS-CSx* first estimates population specific PRS, eg PRS_*ϕ,EUR*_ and PRS_*ϕ,AF R*_, where PRS_*ϕ,x*_ is the standardised PRS for population *x*. For each *ϕ, PRS-CSx* fits the following linear regression to the target population test data *y*

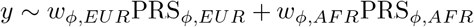

The *ϕ* value and the corresponding regression coefficients for the linear combination of PRS that maximise the coefficient of determination (*R*^2^) in the target population (eg. Africans) test set were used in the validation dataset to calculate the final PRS:

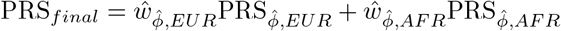

Unlike *BridgePRS, PRS-CSx* does not use European test data to estimate non-European PRS. Therefore, to ensure both methods use the same data GWASs were performed on the European test samples using PLINK v2.0 [23] and then meta-analysed with the GWAS data from the European data METAL [24]. The meta-analysed European GWAS, the GWAS generated from the training samples of the target population, and the LD reference panel generated by the authors of *PRS-CSx* were provided to *PRS-CSx*.

### UK Biobank genotype and sample QC

The UK Biobank (UKB) is a prospective cohort study of around 500,000 individuals recruited across the United Kingdom during 2006-2010. The genetic data is comprised of 488,377 samples genotyped at 805,426 SNPs. Population ancestries were defined by 4-means clustering performed on the first two Principal Components (PCs) of the genotype data. The ancestry of each cluster was defined by the country of birth (field ID: 20115) of the majority of individuals in the cluster. Standard quality control (QC) procedures were then performed on each ancestry cluster independently, any SNP with minor allele frequency <0.01, genotype missingness *>*0.02 or Hardy Weinberg Equilibrium Test *P* -value < 10^−8^ was removed. Samples with high levels of missingness or heterozygosity, with mismatching genetic-inferred and self-reported sex, or with aneuploidy of the sex chromosomes were removed as recommended by the UKB data processing team. A greedy algorithm [25] was used to remove related individuals, with kinship coefficient *>* 0.044, in a way that maximised sample retention. In total, 557,369 SNPs and 387,392 individuals were retained for analysis.

### Imputation

Imputed variants were extracted from imputed UK Biobank data using PLINK v2.0, converting the imputed data into hard-coded genotypes and retaining variants with the following filters: biallelic variants (--max-alleles 2), minor allele frequency greater than 0.001 (–maf 0.001), genotype missingness less than 1% (--geno 0.01) and a MACH info score greater than 0.8 (--mach-r2-filter 0.8).

### Trait selection

We extracted all continuous traits from unique samples in the UK Biobank and performed basic filtering, discarding samples with phenotypic values 6 standard deviation away from the mean. Traits with more than 2,000 samples of African ancestry were extracted. For each trait 300,000 European samples were extracted (retaining at least 10,000 samples for test and validation for each trait) and GWASs run on the genotype data using PLINK v2.0 using --glm. Sex (field ID: 31), age (field ID: 21003), genotyping batch, UK Biobank assessment centre (field ID: 54) and 40 PCs were included as covariates, with fasting time (field ID: 74) and dilution factor (field ID: 30897) also included for blood biochemical traits. LD Score regression [26] was run on the resultant summary statistics and traits were further filtered discarding those with heritability less than 1%. The remaining traits were ranked according to their heritability, and traits correlated with a more heritable trait (absolute Pearson correlation greater than 0.3) were removed resulting in 27 traits. Results are presented for 19 traits which have an *R*^2^ in Africans of greater than 1% for at least one analysis. The sample sizes for each trait and ancestry are shown in Supplementary Table 1.

### Implementation

European, African and South Asian UKB samples were split into three independent groups: training data to construct the GWAS summary statistics; test data, to select best fitting parameters; and validation data, to calculate out-of-sample predictive accuracy. The proportion of samples allocated to each set were 2/3 training, 1/6 test and 1/6 validation. Each GWAS was run in PLINK v2.0 as described above. East Asian samples were split equally between test and validation.

For each trait analyses were run with imputed variants. GWASs were run separately for the training samples of European, African and South Asian ancestry for each of the 27 traits using PLINK v2.0 as described above. All PRS were calculated using two populations: African PRS used African and European UKB GWAS data, South Asian PRS used South Asian and European UKB GWAS data and East Asian PRS used BBJ and European UKB GWAS.

### Application to Bio*Me*

Bio*Me* BioBank samples were genotyped on the Infinium Global Screening Array v1.0 (GSA) platform. Samples were removed with a population-specific heterozygosity rate of greater than *±*6 standard deviations of population-specific mean, along with a call rate of <95%. In addition, samples were removed for exhibiting persistent discordance between EHR recorded and genetic sex. Variants were removed that had a call rate <95%, a Hardy-Weinburg Equilibrium *P* -value threshold of *p* < 10^−5^ in African-American and European-American ancestry, or *p* < 10^−13^ in Hispanic and South Asian ancestry.

PCA was performed; African, South Asian and East Asian samples were selected by clusters on PC plots corresponding to self-reported ancestry. African samples were selected as those with PC 1> 0.0075, PC 2 < -0.0005 and PC 3 > -0.002. South Asian samples were selected as those with -0.01 < PC 3 < -0.004, -0.003 < PC 4 < 0.001 and PC 5 < -0.015. East Asian samples were selected as those with PC 3 < -0.01, PC 4 > 0.001, PC 5 > -0.005 and PC 6 > -0.0035. Supplementary Fig. 6-8 plot the top 6 PCs, with samples coloured by self-reported ancestry and show the thresholds used to select African, South Asian and East Asian ancestry samples.

Imputation was performed using *IMPUTE2* [27] with the 1000G Phase3 v5 reference panel [11]. Variants were first filtered by info score > 0.3. Genotype data for the calculation of PRS in unique individuals was generated for in each of the two ancestry groups separately by first removing variants with minor allele frequency < 1% in the respective Bio*Me* population and then removing one of each pair of variants with duplicate genomic position. Bio*Me* variants were mapped onto the UKB PRS by genomic position (build 37). Variants were coded by their expected allele count (dosage) for the calculation of PRS. Samples with phenotypic values 3 standard deviation away from the mean were excluded.

### Measure of PRS accuracy

Variance explained was calculated as

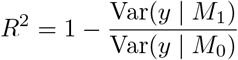

where *M*_*i*_ is the regression model with the PRS (*i* = 1) and without (*i* = 0) and with both models including covariates for top 40 PCs, age, sex, centre and batch, fasting and dilution for the biochemical traits. Variance explained in the applications to Bio*Me* included covariates for age, sex and the top 32 PCs. Standard errors and confidence intervals were calculated by bootstrapping in the R package *boot* [28] using 10,000 replicates.

