## Supplementary Information for "*BridgePRS*: A powerful trans-ancestry Polygenic Risk Score method"

### Reformulation of F-test

Without loss of generality, assume zero centred normally distributed trait data $y$ with variance $\sigma^2$. A linear regression is fit to this data with $n \times k$ covariate matrix $X$ resulting in a maximum likelihood estimates $\hat{\beta}$. The $F$ statistic is defined by the residual sum of squares of the null and alternative models ($\mathrm{RSS}_0$ and $\mathrm{RSS}_1$) as follows:

$$
\begin{aligned}
F &= \frac{n-k}{k}\left(\frac{\mathrm{RSS}_0 - \mathrm{RSS}_1}{\mathrm{RSS}_1}\right) \\
&= \frac{n-k}{k}\left(\frac{y^T y - (y - X\hat{\beta})^T (y - X\hat{\beta})}{(y - X\hat{\beta})^T (y - X\hat{\beta})}\right) \\
&= \frac{n-k}{k}\left(\frac{n\sigma^2}{(y - X\hat{\beta})^T (y - X\hat{\beta})} - 1\right)
\end{aligned}
$$

$$= \frac{n-k}{k}\left(\frac{n\sigma^2}{y^Ty - \hat{\beta}^TX^Ty - y^TX\hat{\beta} + \hat{\beta}^TX^TX\hat{\beta}} - 1\right)$$

since $\hat{\beta} = (X^TX)^{-1}X^Ty$

$$= \frac{n-k}{k}\left(\frac{n\sigma^2}{n\sigma^2 - \hat{\beta}^TX^TX\hat{\beta}} - 1\right)$$

$$= \frac{n-k}{k}\left(\frac{\hat{\beta}^TX^TX\hat{\beta}}{n\sigma^2 - \hat{\beta}^TX^TX\hat{\beta}}\right)$$

$$= \frac{n-k}{kn\sigma^2}\hat{\beta}^TX^TX\hat{\beta}\left(1 - \frac{1}{n\sigma^2}\hat{\beta}^TX^TX\hat{\beta}\right)^{-1}$$

$\frac{\beta^TX^TX\beta}{\sigma^2}$ is the variance explained by the locus, therefore, assuminhg this is small a first order Taylor approximation can be used to give

$$\approx \frac{n-k}{kn\sigma^2}\hat{\beta}^TX^TX\hat{\beta}$$

| | European | | African | | | South Asian | | East Asian | |
|---|---|---|---|---|---|---|---|---|---|
| | Train | Train | Test | Valid. | Train | Test | Valid. | Test | Valid. |
| Standing height | 257675 | 4862 | 1215 | 1214 | 6666 | 1666 | 1665 | 1235 | 1235 |
| Body mass index (BMI) | 257327 | 4853 | 1213 | 1212 | 6653 | 1662 | 1662 | 1234 | 1233 |
| Platelet count | 250404 | 4657 | 1163 | 1163 | 6538 | 1634 | 1633 | 1204 | 1203 |
| C-reactive protein | 176422 | 3281 | 820 | 819 | 4581 | 1145 | 1144 | 1034 | 1034 |
| Neutrophill count | 249945 | 4642 | 1160 | 1159 | 6515 | 1628 | 1627 | 1202 | 1201 |
| Mean corpuscular volume | 250359 | 4660 | 1164 | 1164 | 6541 | 1634 | 1634 | 1204 | 1203 |
| Apolipoprotein A | 161594 | 3029 | 756 | 756 | 4182 | 1045 | 1044 | - | - |
| Reticulocyte percentage | 246092 | 4531 | 1132 | 1131 | 6367 | 1591 | 1590 | - | - |
| Alkaline phosphatase | 177485 | 3293 | 823 | 822 | 4611 | 1152 | 1152 | 1034 | 1034 |
| LDL direct | 177383 | 3291 | 822 | 821 | 4603 | 1150 | 1150 | 1034 | 1034 |
| Monocyte count | 249782 | 4639 | 1159 | 1158 | 6503 | 1625 | 1625 | 1202 | 1202 |
| RBC distribution width | 249871 | 4645 | 1161 | 1160 | 6523 | 1630 | 1630 | - | - |
| Urea | 177343 | 3289 | 821 | 821 | 4603 | 1150 | 1149 | - | |
| Triglycerides | 177317 | 3289 | 821 | 821 | 4603 | 1150 | 1149 | 1034 | 1034 |
| Basophill % | 249368 | 4623 | 1155 | 1154 | 6495 | 1623 | 1622 | 1199 | 1199 |
| Total protein | 162440 | 3045 | 761 | 760 | 4190 | 1047 | 1046 | 1034 | 1034 |
| Sodium in urine | 250253 | 4734 | 1183 | 1182 | 6497 | 1624 | 1623 | - | - |
| IGF-1 | 176643 | 3272 | 817 | 817 | 4583 | 1145 | 1144 | - | - |
| Eosinophill count | 249594 | 4629 | 1157 | 1156 | 6503 | 1625 | 1624 | 1198 | 1197 |

**Table 1** UK Biobank train, test and validation sample sizes for each ancestry group across the 19 traits analysed. Across all traits 10,000 samples were used as test data. Numbers for East Asians are only shown for traits overlapping with those with available BBJ summary statistics.

| | African | South Asian | East Asian |
|---|---|---|---|
| Standing height | 4061 | 560 | 589 |
| Body mass index | 4084 | 560 | 595 |
| LDL direct | 1998 | 217 | 220 |
| Mean corpuscular volume | 2412 | 360 | 366 |
| Platelet count | 2459 | 368 | 369 |
| Monocyte count | 2229 | 283 | 310 |
| Neutrophill count | 2226 | 284 | 309 |
| Eosinophill count | 2204 | 268 | 306 |
| Red blood cell distribution width | 2398 | 366 | - |

**Table 2** Bio*Me* Biobank sample sizes for individuals of African, South Asian ans East Asian ancestry.

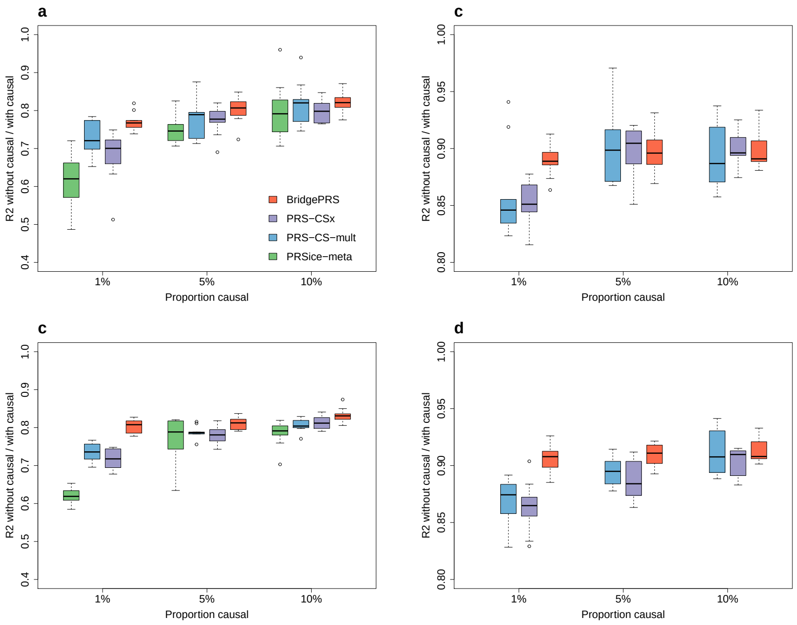


**Fig. 1** Relative loss in removing causal variants from analysis measured by ratio of models' phenotypic variance explained ($R^2$) without and with the causal variants included for *BridgePRS*, *PRS-CSx*, *PRS-CS-mult* and *PRSice-meta* across six siumulation scenarios, ten simulated phenotypes per scenerio. **a** African ancestry samples for phenotypes with $h^2 = 0.25$. **b** East Asian ancestry samples for phenotypes with $h^2 = 0.25$. **c** African ancestry samples for phenotypes with $h^2 = 0.5$. **d** East Asian ancestry samples for phenotypes with $h^2 = 0.5$. *PRSice-meta* results for East Asian analyses were unstable and removed for clarity.

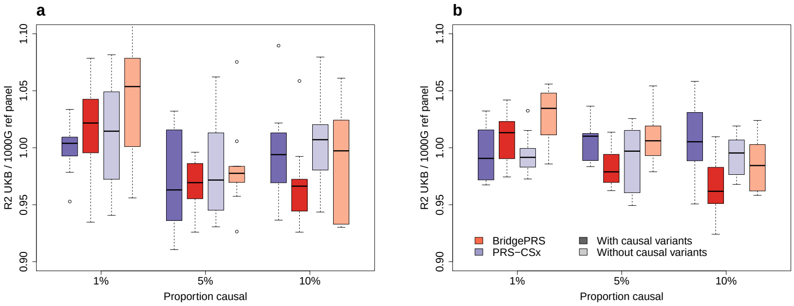


**Fig. 2** Ratio of phenotypic variance explained ($R^2$) comparing models fit using UKB and 1000G LD reference panels for *BridgePRS* and *PRS-CSx* across six siumulation scenarios, ten simulated phenotypes per scenerio. **a** African ancestry samples for phenotypes with $h^2 = 0.25$. **b** East Asian ancestry samples for phenotypes with $h^2 = 0.25$.

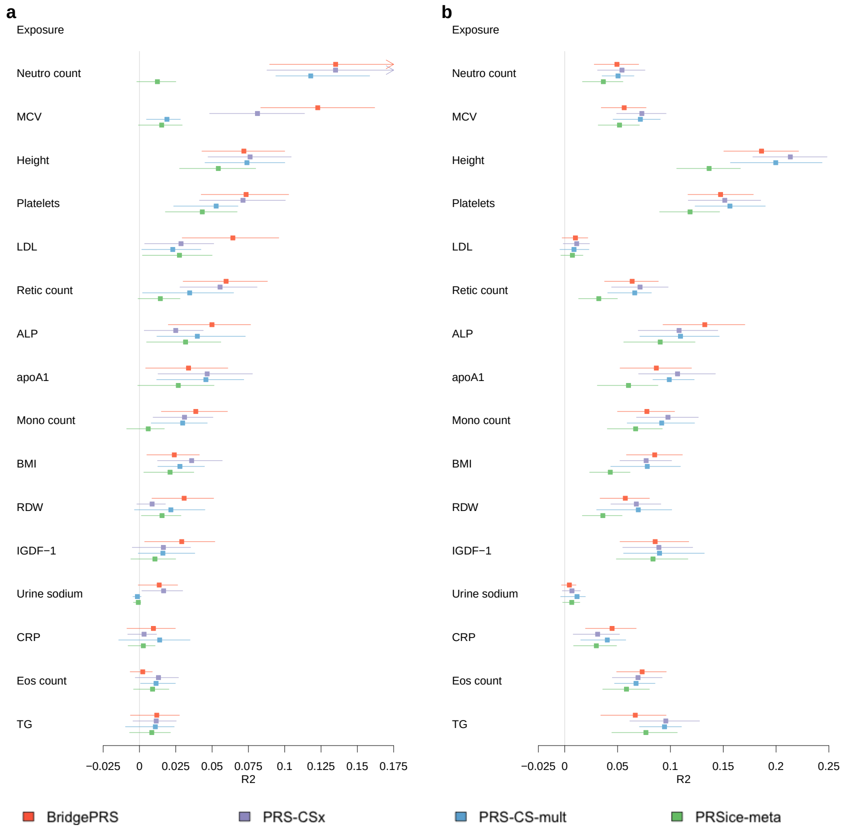


**Fig. 3** Phenotypic variance explained ($R^2$ point estimates and 95% confidence intervals) by *BridgePRS*, *PRS-CSx*, *PRS-CS-mult* and *PRSice-meta* in samples of African and South Asian ancestry in the UK Biobank. **a** African ancestry samples. **b** South Asian ancestry samples. Neutro count=Neutrophill count, MCV=Mean corpuscular volume, Platelets=Platelet count, Retic count=Reticulocyte percentage, ALP=Alkaline phosphatase, Mono count=Monocyte count, apoA1=Apolipoprotein A, BMI=Body mass index, RDW=Red blood cell distribution width, Eos count=Eosinophill count, TG=Triglycerides, Baso %=Basophill percentage, CRP=C-reactive protein

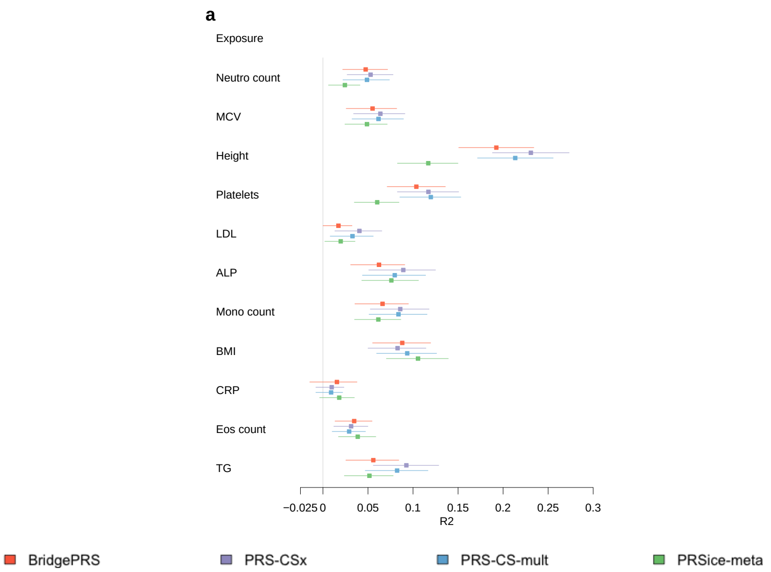


**Fig. 4** Phenotypic variance explained ($R^2$ point estimates and 95% confidence intervals) by *BridgePRS*, *PRS-CSx*, *PRS-CS-mult* and *PRSice-meta* in samples of East Asian ancestry in the UK Biobank. Neutro count=Neutrophill count, MCV=Mean corpuscular volume, Platelets=Platelet count, ALP=Alkaline phosphatase, Mono count=Monocyte count, BMI=Body mass index, Eos count=Eosinophill count, TG=Triglycerides

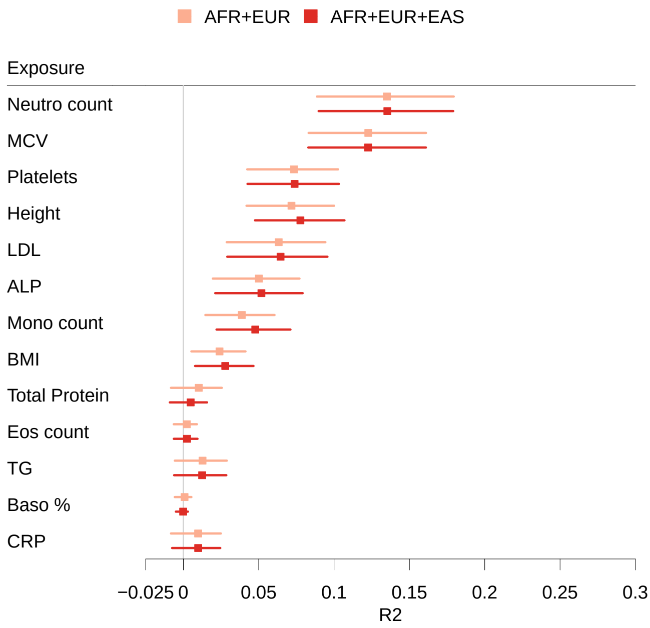


**Fig. 5** Comparison of phenotypic variance explained ($R^2$ point estimates and 95% confidence intervals) by *BridgePRS* in samples of African ancestry in the UK Biobank training PRS using (1) AFR+EUR UKB summaray statistic data and (2) AFR+EUR UKB +EAS BBJ summaray statistic data.

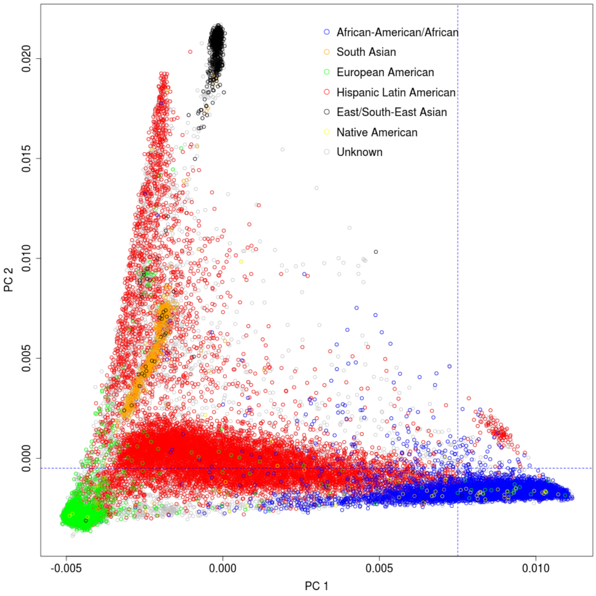


**Fig. 6** Plot of PC 1 v PC 2 for Bio*Me* samples. Samples are coloured by self reported ancestry. Samples in bottom right corner indicated by blue dashed line were defined as African ancestry in our analyses.

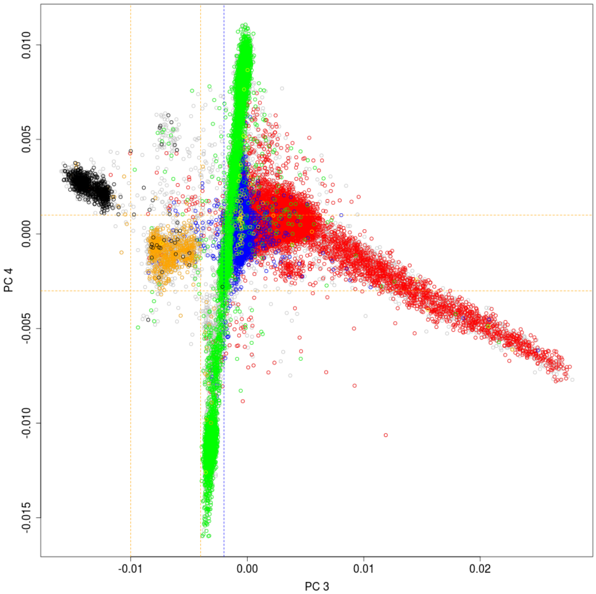


**Fig. 7** Plot of PC 3 v PC 4 for Bio*Me* samples. Samples are coloured by self reported ancestry. Samples in box defined by orange lines were defined as South Asian ancestry in our analyses. Samples to left of blue vertical were excluded from African analyses.

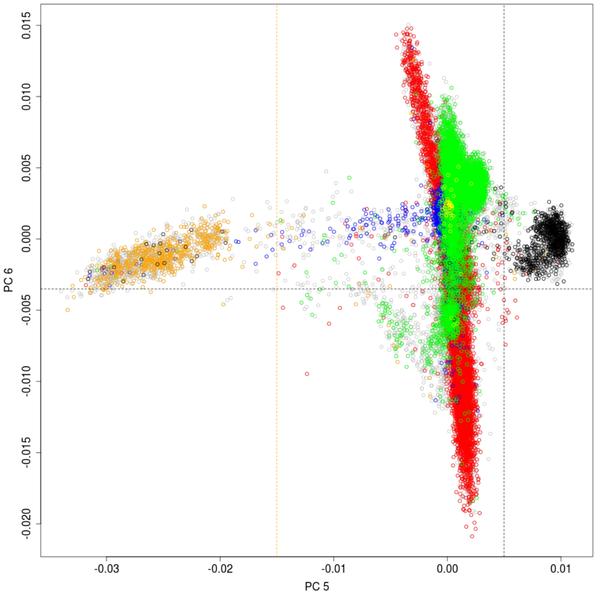


**Fig. 8** Plot of PC 5 v PC 6 for Bio*Me* samples. Samples are coloured by self reported ancestry. Samples to right of orange vertical were excluded from South Asian analyses.
